# A direct glia-to-neuron natural transdifferentiation ensures nimble sensory-motor coordination of male mating behaviour

**DOI:** 10.1101/285320

**Authors:** Laura Molina-García, Byunghyuk Kim, Steven J. Cook, Rachel Bonnington, Jack M. O’Shea, Michele Sammut, Sophie P. R. Gilbert, David J. Elliott, David H. Hall, Scott W. Emmons, Arantza Barrios, Richard J. Poole

**Affiliations:** Department of Cell and Developmental Biology, University College London, London WC1E 6BT, UK; Department of Genetics, Albert Einstein College of Medicine, Bronx, New York 10461, USA; Department of Life Science, Dongguk University-Seoul, Goyang 10326, Republic of Korea; Dominick P. Purpura Department of Neuroscience, Albert Einstein College of Medicine, Bronx, New York 10461, USA; Department of Biological Sciences, Columbia University, New York, NY 10027; Centre for Discovery Brain Sciences, Edinburgh University, Edinburgh EH8 9XD, UK

## Abstract

Sexually dimorphic behaviours require underlying differences in the nervous system between males and females. The extent to which nervous systems are sexually dimorphic and the cellular and molecular mechanisms that regulate these differences are only beginning to be understood. We reveal here a novel mechanism to generate male-specific neurons in *Caenorhabditis elegans*, through the direct transdifferentiation of sex-shared glial cells. This glia-to-neuron cell fate switch occurs during male sexual maturation under the cell-autonomous control of the sex-determination pathway. We show that the neurons generated are cholinergic, peptidergic and ciliated putative proprioceptors which integrate into male-specific circuits for copulation. These neurons ensure coordinated backward movement along the mate’s body during mating. One step of the mating sequence regulated by these neurons is an alternative readjustment movement performed when intromission becomes difficult to achieve. Our findings reveal programmed transdifferentiation as a developmental mechanism underlying flexibility in innate behaviour.

## Introduction

The coordinated execution of innate, stereotyped sexual behaviours, such as courtship and mating, requires sexually dimorphic sensory-motor circuits that are genetically specified during development (reviewed in ^1–3^). Studies in the nematode *Caenorhabditis elegans*, in which the development and function of neural circuits can be interrogated with single cell resolution, have revealed two general developmental mechanisms underlying sexual dimorphism in the nervous system. The first involves the acquisition of sexually dimorphic features in sex-shared neurons during sexual maturation, which include changes in terminal gene expression, such as odorant receptors, neurotransmitters and synaptic regulators ^4-9 10,11^. The second mechanism involves the generation of sex-specific neurons ^12–14^. Sex-specific neurons are primarily involved in controlling distinct aspects of reproductive behaviours, such as egg-laying in the hermaphrodite and mating in the male (reviewed in ^15^). Generation of sex-specific neurons requires sex-specific cell death ^16^ or neurogenesis events resulting from sex differences in the cell division patterns and neurodevelopmental programmes of post-embryonic cell lineages (reviewed in ^3^). Here we identify a third, novel way to generate sexual dimorphism in the nervous system.

In one of his seminal papers, John Sulston described a sexual dimorphism in the phasmid sensilla of adult animals ^13^. The phasmid sensillum is one of the seven classes of sense organs that are common to both sexes in *C. elegans*. These sense organs are organised in sensilla which are concentrated in the head and the tail ^17-19 20^. Each sensillum is composed of the dendrites of one or more sensory neurons enveloped by a channel, usually composed of a single sheath glial cell and a single socket glial cell. These sensilla can be viewed as part of an epithelium, continuous with the skin, and are shaped by mechanisms shared with other epithelia ^21^. Socket glial cells are highly polarised and adhere to the hypodermis at the distal end of their process where they form a small, ring-like hollow pore in the cuticle through which the sensory dendrites can access the outside world. The bilateral phasmid sensilla (Fig.1), situated in the tail, are unusual in that they are each composed of two socket glial cells (PHso1 and PHso2). John Sulston observed that in juvenile animals (L2 stage) of both sexes, PHso1 forms the primary pore (^13^; Fig.1A). At adulthood, the hermaphrodite retains a similar structure (Fig.1B). In males, however, it is PHso2 that forms the main pore and PHso1 was described as having retracted from the hypodermis and to protrude into the phasmid sheath (Fig.1C). It was also described to contain basal bodies, a structural component of cilia, in the region enveloped by the phasmid sheath. As sensory neurons are the only ciliated cells in *C. elegans* ^22^, this is suggestive of neuronal fate, yet Sulston observed no other neuronal characteristics ^13^. Because we have previously shown that in the amphid sensillum (a similar organ located in the head), the amphid socket glial cell (AMso) acts as a male-specific neural progenitor that, during sexual maturation, divides to self-renew and generate the MCM neurons ^14^, we sought to investigate the PHso1 cells in more detail.

**Figure 1:**
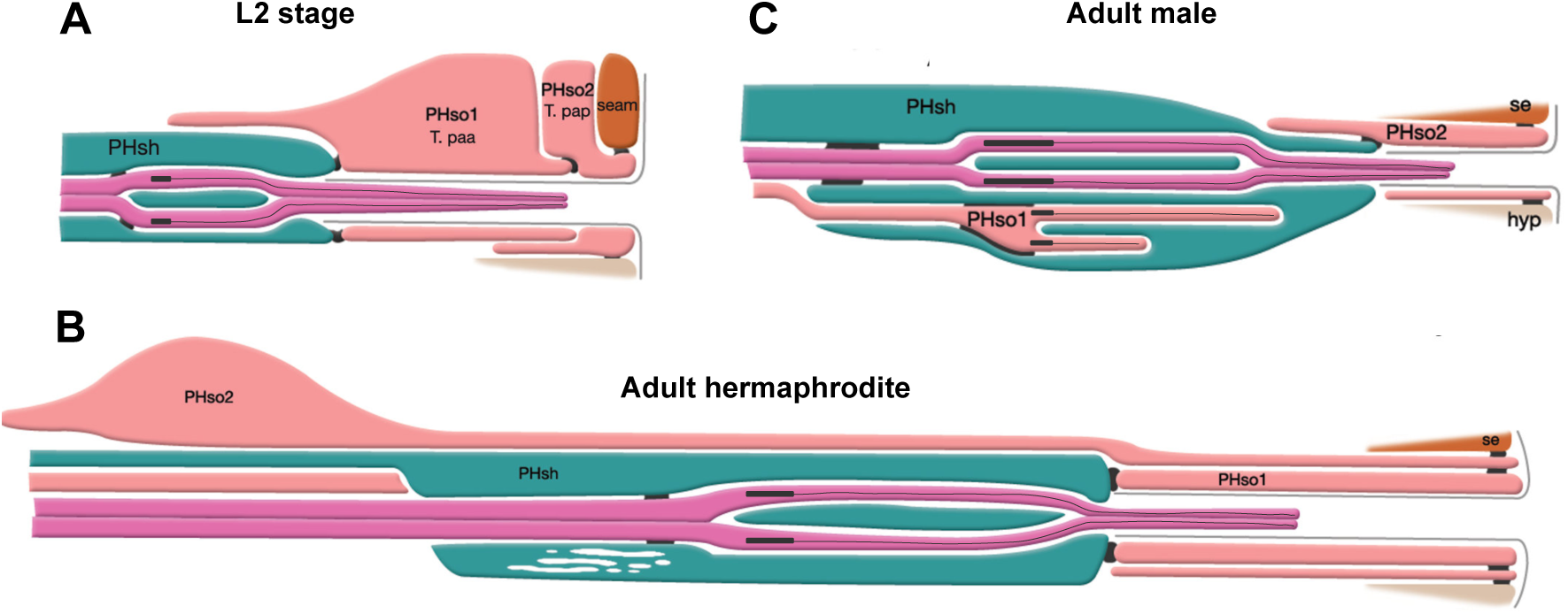
The phasmid sensillum. Diagram of the phasmid sensillum in either sex at the L2 larval stage (A), in adult hermaphrodites (B) and in adult males (C). The socket glial cells (PHso1 and PHso2) are coloured in light pink; the sheath glial cells (PHsh) in green; and the ciliated dendrites of the phasmid sensory neurons, in pink. The adherens junctions are depicted as black lines between cells. Axonemes and cilia are marked as black bars and black lines inside the dendrite tips. Each phasmid opens to the exterior on the extreme right (posterior), where grey lines mark the cuticle borders of the phasmid pore and fan. Hypodermis (hyp), seam (se). Diagram has been modified from and is used with permission from http://www.wormatlas.org

We find that during sexual maturation (L4 stage), the sex-shared PHso1 glial cells acquire sexually dimorphic function by undergoing a direct glia-to-neuron transdifferentiation that results in the production of male-specific neurons. This plasticity is regulated cell-intrinsically by the sex-determination pathway. These previously unnoticed neurons, which we term PHDs, are putative proprioceptors that regulate male locomotion during specific steps of mating. One of these steps is a novel readjustment movement performed when intromission becomes difficult to achieve. Our results reveal sex-specific direct transdifferentiation as a novel mechanism for generating sex-specific neurons and also show the importance of proprioceptive feedback during the complex steps of mating for successful reproduction.

## Results

### The sex-shared PHso1 cells undergo glia-to-neuron transdifferentiation in males

Using a *lin-48/OVO1* reporter transgene to identify and visualise the PHso1 cell bodies both before and during sexual maturation ^23,24^, we observed no distinguishable differences between the hermaphrodite and male PHso1 cells at the L3 stage (Fig. 2A). The PHso1 cells display a polarised morphology and a visible socket structure in both sexes. In hermaphrodites, this morphology is maintained during the transition to adulthood and PHso1 cells elongate as the animal grows. In males, in contrast, PHso1 displays morphological changes that can be observed during the L4 stage, the last larval stage before adulthood (Fig. 2A). We find that in early L4 PHso1 retracts its socket process and extends a short dendrite-like posterior projection into the PHsh cell and a long axon-like anterior process projecting towards the pre-anal ganglion region. A time-series of carefully staged animals reveals that axon-like outgrowth is complete by the time of tail-tip retraction (Fig. 2B). These observations corroborate those of John Sulston and identify the stage of sexual maturation as the time during which the PHso1 cells undergo radical remodelling in males. This remodelling involves quite a remarkable change in morphology from a socket-glial morphology to a neuron-like morphology. Importantly, we do not observe any cell division of the PHso1 cells during this process. We only ever observe one bilateral pair of cells expressing *lin-48* and only two bilateral pairs of cells (PHso1 and PHso2) expressing the glial subtype marker *grl-2* during the change in morphology (Fig. 2 and Fig. 3). Together, these observations indicate that PHso1 cells undergo direct remodelling during which they may directly acquire neuronal fate.

**Figure 2.**
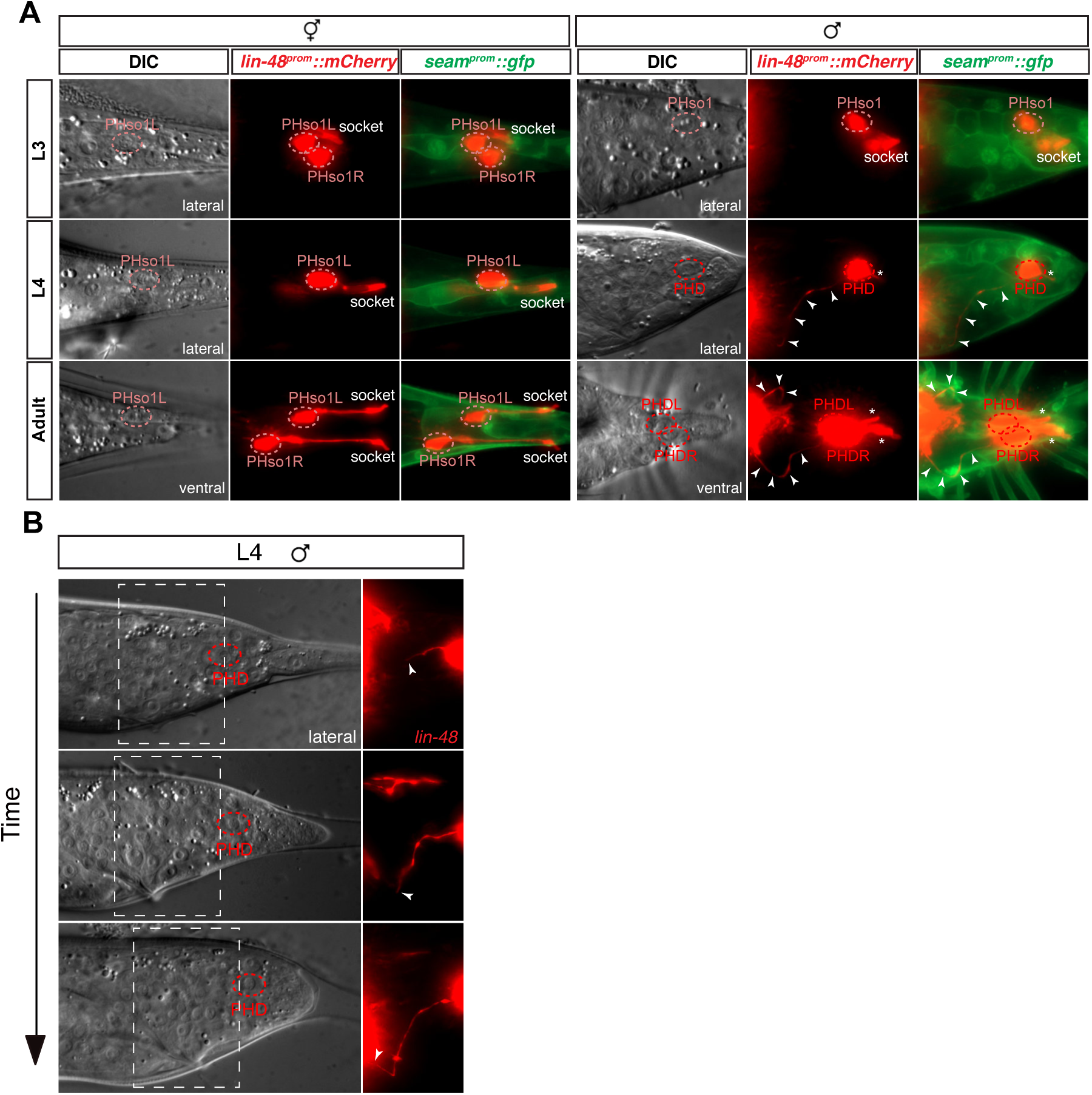
The sex-shared PHso1 cells undergo glia-to-neuron morphological changes in males. A.Expression of *lin-48::mCherry* and the *seam::gfp* (*wrt-2::gfp)* reporter transgenes in PHso1 of hermaphrodites (left panel) and males (right panel) at the third (L3) and fourth (L4) larval stages and in adults. The images show the morphological transformation of male PHso1 into the PHD neuron during sexual maturation. Arrowheads label the process extending from the PHD into the pre-anal ganglion. Starts indicate the dendritic process of PHD. B. DIC and fluorescent images of fourth-larval stage (L4) male tails at early, mid and late stages. The outgrowth of the anterior process of the PHso1/PHD cell can be seen by expression of a *lin-48::tdTomato* transgene.

**Figure 3:**
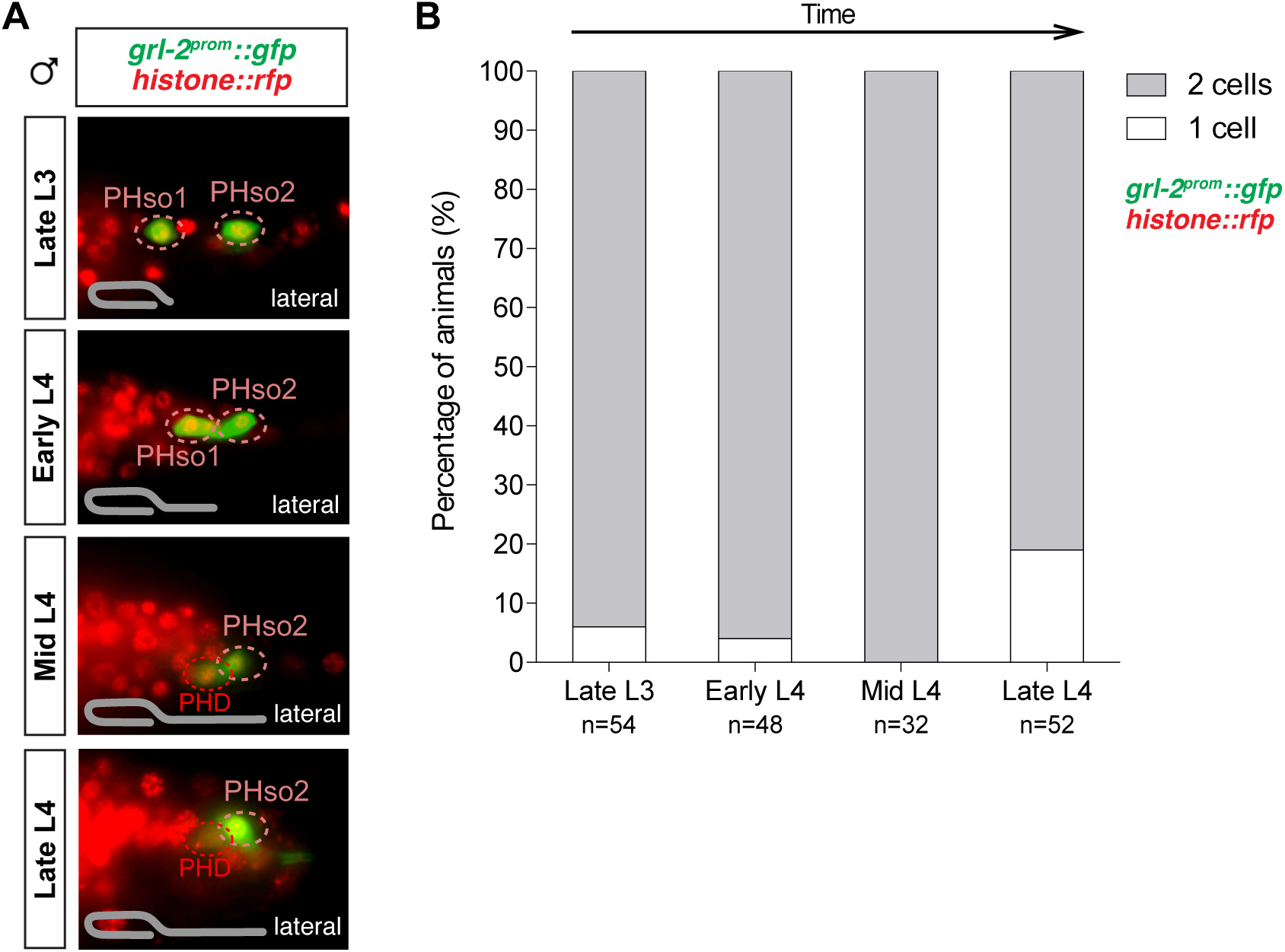
The PHso1 cells do not divide during sexual maturation. In addition to observing only one *lin-48* expressing cell (PHso1) per side, only two *grl-2* expressing cells (PHso1 and PHso2) per side are observed at any time during the glia-to-neuron remodelling of PHso1. Chromosome condensation (as visualised with a *histone::rfp* reporter) was never observed in either *grl-2* expressing cell. A range of carefully staged animals were scored before, during and after remodelling. A. Fluorescent images of male tails containing both a *grl-2::gfp* transgene and a *histone::rfp* transgene at different stages of development. A range of animals was scored and subdivided into four categories based on gonad migration and tail morphogenesis: late L3 - from the stage at which the gonad begins its posterior migration to the point at which the gonad crosses itself (~28-32 hrs); early L4 - from the stage at which the gonad crosses itself to 50% extension towards the tail (~32-34 hours); mid L4 - from 50% extension of the gonad towards the tail to 100% extension (~34-36 hours); late L4 - from the start of to mid tail tip retraction. The gonad morphology for the oldest animals in each set is illustrated in grey. B. Quantification of the number of cells expressing *grl-2* (and displaying uncondensed chromosomes) per side at each of the above stages.

To determine whether the PHso1 cells acquire neuronal characteristics at the level of gene expression in addition to morphology, we analysed and compared the expression of glial and neuronal markers in the PHso1 and PHso2 cells of males and hermaphrodites. We find that both cells express the panglial microRNA *mir-228* ^25^ and the AMso/PHso glial subtype marker *grl-2* (a hedgehog-like and Ground-like gene protein ^26^) at equivalent levels in adult hermaphrodites (Fig. 4A). In males, in contrast, *mir-228* and *grl-2* expression is noticeably dimmer in PHso1 than in PHso2 by the mid-to-late L4 stage and completely absent by day two of adulthood (Fig. 4A, B and C). Importantly, expression of these genes is equal in brightness in both cells at the late L3 and early L4 stages (Fig. 4B and C). By the L4 moult, PHso1 begins to express the pan-neuronal marker *rab-3* (a synaptic vesicle associated Ras GTPase ^27^) in males but not in hermaphrodites, and this expression persists throughout adulthood (Fig. 4B and C). In addition to *rab-3* expression we also observe expression of other neuronal markers such as: the vesicle acetylcholine transporter *unc-17* ^28,29^ (Fig. 4D); the phogrin orthologue for dense-core vesicle secretion *ida-1* ^30^ (Fig. 4E); the intraflagellar transport component required for proper sensory cilium structure *osm-6* ^31^ (Fig. 4F); and the IG domain containing protein *oig-8* ^32^ (Fig. 6A). None of these neuronal markers are observed in PHso2 or in the hermaphrodite PHso1. The switch in gene expression in PHso1 is therefore initiated concomitantly with morphological changes during sexual maturation and in a male-specific manner. Together, these data demonstrate that the male PHso1 glial cells transdifferentiate into a novel class of ciliated neurons which we have termed phasmid D neurons (PHDs).

**Figure 4:**
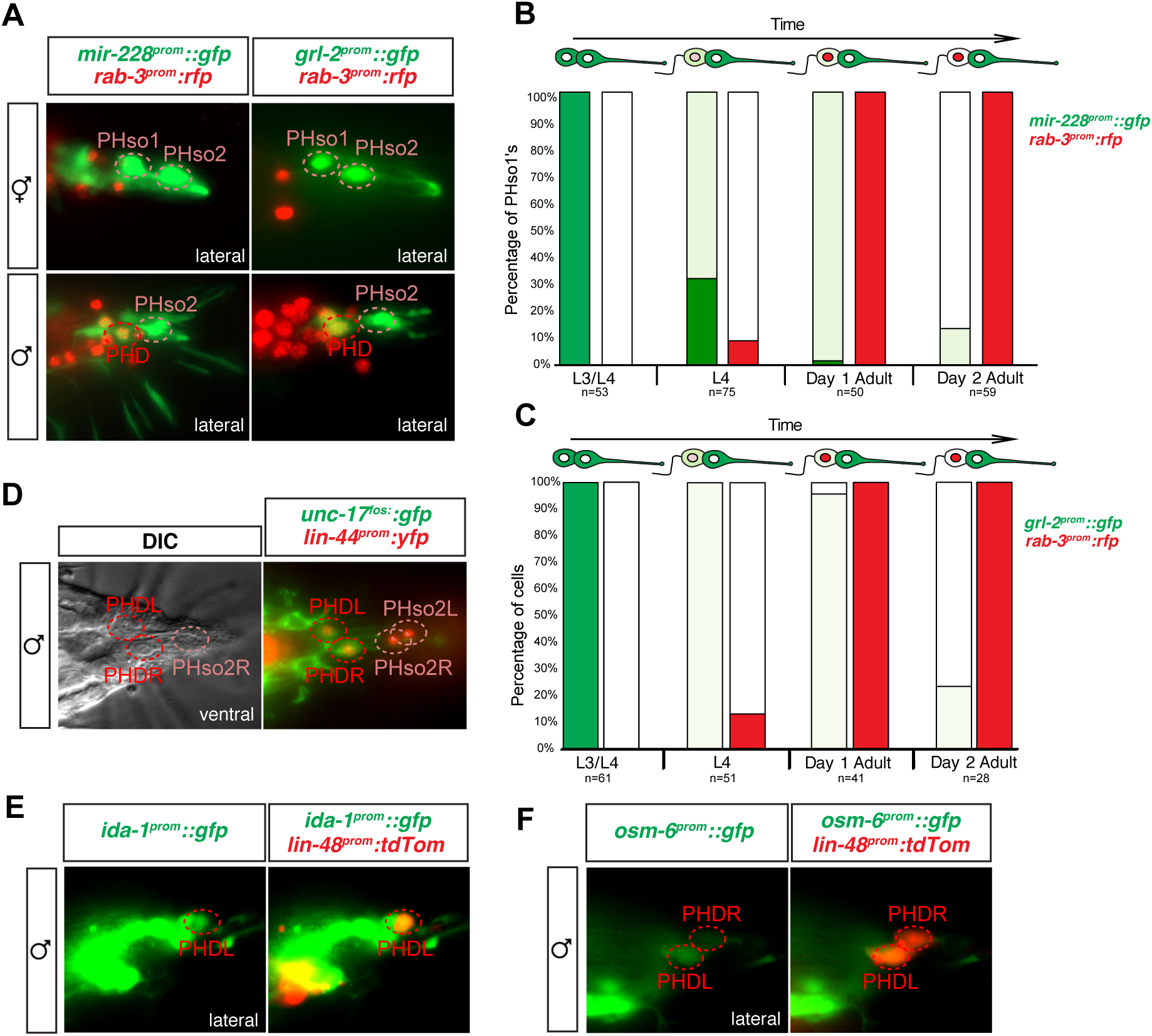
The sex-shared PHso1 cells undergo glia-to-neuron molecular changes in males. A. Expression of the glial marker reporter transgenes *mir-228::gfp* and *grl-2::gfp* and pan-neuronal marker *rab-3::RFP* in the phasmid socket cells (PHso1 and PHso2) and the PHD neuron in adult animals. B. Bar chart showing the percentage of PHso1/PHD cells expressing the pan-glial marker reporter transgene *mir-228::gfp* and the pan-neuronal marker reporter transgene *rab-3::rfp* scored concomitantly in males at different stages of development and at adulthood. Intensity of the *mir-228::gfp* reporter transgene in the PHso1/PHD cells was assessed in comparison with PHso2: dark green indicates PHso1/PHD = PHso2 and light green indicates PHso1/PHD < PHso2. C. As B but for the subtype-specific glial marker *grl-2::gfp* scored concomitantly with *rab-3::rfp*. D. Expression of the acetylcholine vesicle uploader reporter transgene *unc-17::gfp* in the PHD neurons of adult males. The *lin-44::yfp* reporter transgene has been coloured red. E. Expression of an *osm-6::gfp* reporter transgene in the PHD neurons of adult males, which are co-labelled with a *lin-48::tdTomato* transgene. F. Expression of an *ida-1::gfp* reporter transgene in the PHD neurons of adult males, which are co-labelled with a *lin-48::tdTomato* transgene.

### Biological sex regulates PHso1-to-PHD transdifferentiation cell-autonomously

We next addressed whether the PHso1-to-PHD cell fate switch is regulated by the genetic sex of the cell rather than the sex of the rest of the animal, in a manner similar to that of several other sexual dimorphisms in *C. elegans* ^5-7,10,14,33-36^. To uncouple the sex of PHso1 from the rest of the animal, we drove expression of *fem-3* in PHso1 (and PHso2) under the *grl-2* promoter. *fem-3* inhibits the expression of *tra-1*, a downstream target of the sex-determination pathway that activates hermaphrodite development and inhibits male development ^37^ (and reviewed in ^38^). Therefore, cell-specific expression of a *fem-3* transgene will masculinise a cell in an otherwise hermaphrodite background. We find that *fem-3* expression specifically in the PHso1 transforms it into a PHD-like neuron, resulting in the upregulation of *ida-1* and *rab-3* expression and the acquisition of neuronal morphology (Fig. 5). This indicates that the competence of PHso1 to transdifferentiate into PHD is cell-intrinsic and based on genetic sex.

**Figure 5:**
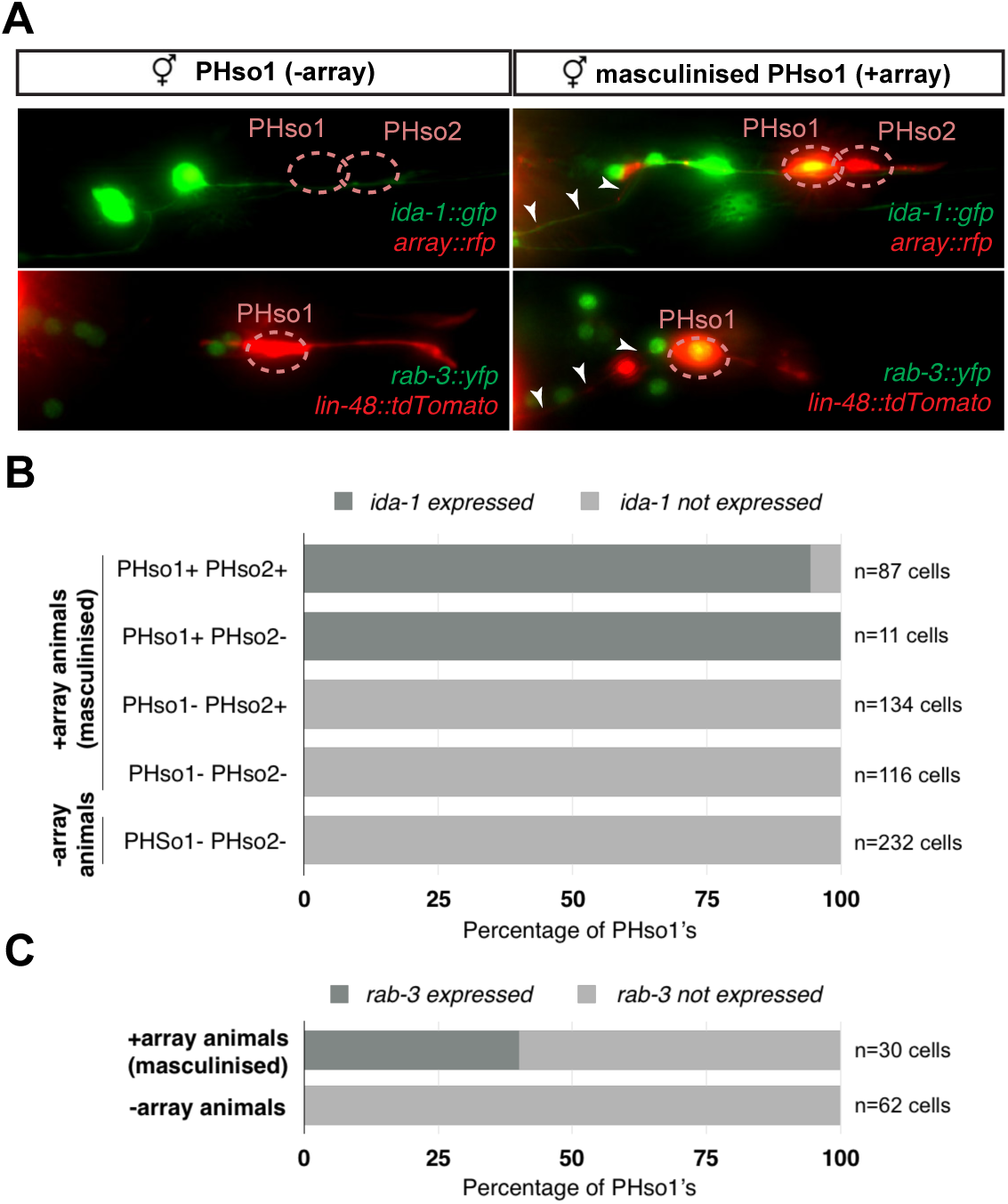
PHso1-to-PHD plasticity is intrinsically regulated. **A.** Expression of the *ida-1::gfp* and *rab-3::yfp* reporter transgenes in adult hermaphrodites carrying the masculinising array *grl-2::fem-3::mCherry* in PHso1 (right panel) and in non-array carrying hermaphrodites (left panel). **G.** Bar chart showing the percentage of PHso1 and PHso2 cells expressing the *ida-1::gfp* reporter transgenes in adult hermaphrodites carrying or not carrying the masculinising *grl-2::fem-3::mCherry* array. **H.** Bar chart showing the percentage of PHso1 cells expressing the *rab-3::yfp* reporter transgene in adult hermaphrodites carrying or not carrying the masculinising *grl-2::fem-3::mCherry* array.

### PHDs are sensory neurons of male-specific copulation circuits

To further establish the neuronal characteristics of PHDs, we examined their synapses and ultrastructure, and mapped their full wiring diagram. We identified a ~1kb promoter region that specifically drives the expression of *oig-8* in the PHD neurons. Expression of a *rab-3* translational fusion under the control of this *oig-8* promoter reveals that PHDs form synapses in the pre-anal ganglion, where the synaptic-vesicle associated mCherry-tagged RAB-3 protein can be observed (Fig. 6A). At the ultrastructural level, we observed both synaptic and dense core vesicles in PHD, proximal to the cell body (Fig. 6B). Ultrastructural analysis of the PHD dendrite in seven different animals independently confirms John Sulston’s original observation on the presence of cilia, including the basal body and axonemes. The basal body of PHD lies just dorsal and anterior to the phasmid sheath channel containing the PHA and PHB cilia, with 2, 1 or zero short cilia extending within the phasmid sheath very close to PHA and PHB cilia, but within a separate sheath channel. Interestingly, the PHD cilia can lie in variable positions relative to PHA and PHB cilia, medial, lateral, dorsal or ventral. In addition, we observe that the dendrite is more elaborate than previously described and has a number of unciliated finger-like villi within the phasmid sheath cell, proximal to the basal body (Fig. 6C and Fig. S1). Through reconstruction of serial electron micrographs, we identified all the synaptic partners of PHD (Table S1). PHD neurons project from the dorsolateral lumbar ganglia, anteriorly and then ventrally along the posterior lumbar commissure and into the pre-anal ganglion. In the pre-anal ganglion, these establish chemical synapses and gap junctions with sensory neurons and interneurons, most of which are male-specific (Fig. 6D). Previously, the axonal process of PHD was attributed to the R8B ray neuron ^4^. A re-examination of the lumbar commissure allowed us to disambiguate PHD’s axon. PHDs receive synaptic input from male-specific ray sensory neurons involved in the initiation of the mating sequence in response to mate contact ^39,40 41^. The main PHD output is to the male-specific EF interneurons (35.4% of chemical synapses), both directly and through their second major post-synaptic target, the sex-shared PVN interneuron. The EF interneurons are GABAergic ^8^ and synapse onto the AVB pre-motor interneurons which drive forward locomotion. Other PHD outputs include the PVV (male-specific) and PDB (sex-shared but highly dimorphic) interneurons, which synapse onto body-wall muscle, and the cholinergic male-specific interneurons PVY and PVX whose output is to the AVA pre-motor interneurons, which drive backward locomotion. PVN, PVY and PVX form disynaptic feed-forward triplet motifs that connect PHD strongly to the locomotion circuit interneurons AVB and AVA. The pattern of connectivity of PHD suggests a possible role in male mating behaviour.

**Figure 6:**
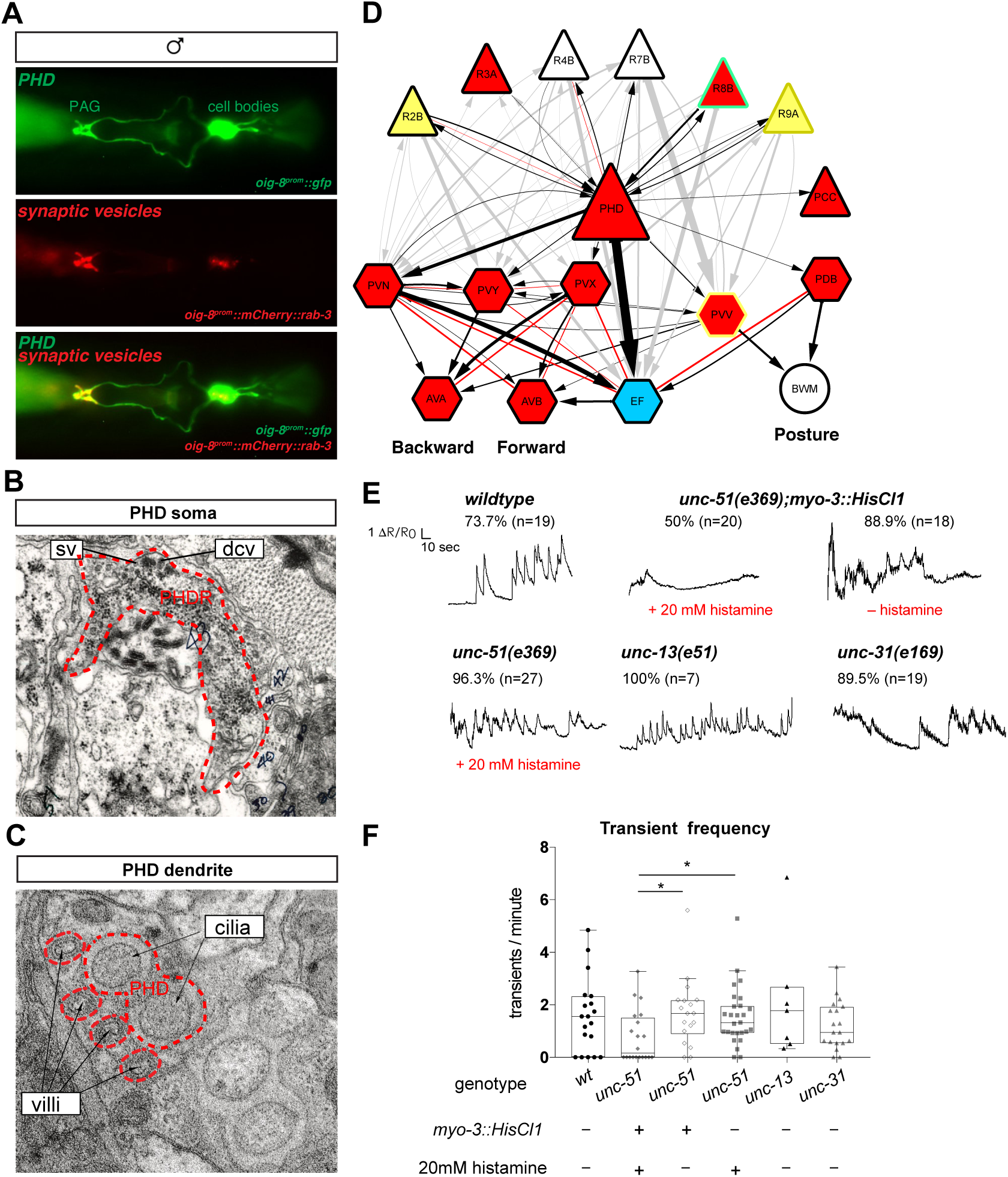
The PHDs are putative proprioceptive neurons of male-specific copulation circuits. A. Expression of the *oig-8::mCherry::rab-3* and *oig-8::gfp* reporter in the PHD neurons of adult males. The synapses made by the PHDs in the pre-anal ganglion (PAG) can be observed (ventral view). B. Electron micrographs of the soma of a PHD neuron of an adult male. sv, synaptic vesicle; dcv, dense core vesicles; PHDR, right PHD neuron. C. As B for PHD dendrite. D. Diagram depicting the connectivity of the PHD neurons with their main pre-synaptic inputs (ray neurons) and post-synaptic targets. The connections between the ray neurons (RnA/B) and their postsynaptic targets independently of PHD are indicated in grey. Arrows and red lines indicate chemical and electrical synaptic connections, respectively. The thickness of the arrows is proportional to the anatomical strength of their connections (# serial sections). Neurons are coloured coded according to their neurotransmitter: red, cholinergic; yellow, glutamatergic; dark yellow, dopamine; blue, GABAergic; green, serotonergic; white, orphan. Note that some neurons (R8B, R9A, PVV) express more than one neurotransmitter. E. Example traces showing PHD activity as normalised GCaMP/RFP fluorescence ratio in restrained animals. Traces are shown for a *wildtype* male, an *unc-51(e359)* mutant with and without a histamine-inducible silencing transgene in muscle (*myo-3::HisCl1*), a mutant in synaptic transmission (*unc-13*) and a mutant in dense core vesicle exocytosis (*unc-31*, CADPS/CAPS). The proportion of traces where calcium peaks were identified is indicated for each genotype and treatment. n= number of neurons imaged F. Plots of frequency values of calcium transients per neuron. Dots represent individual neurons imaged. Tukey box-and-whisker plots indicate the interquartile ranges and median. * *P*<0.05; One-way anova with multiple comparisons. Two groups were compared: *unc-51* genotypes and treatments and another group including *wt, unc-13* and *unc-31.* Only statistically significant comparisons are indicated.

### PHDs are putative proprioceptive neurons

We next sought to establish the function of the PHD neurons. We began by asking what sensory stimuli might activate them. To monitor neuronal activity, we co-expressed GCaMP6f and RFP in the PHDs and performed ratiometric measurements of fluorescence. We imaged calcium transients in restrained animals glued to a slide without anaesthetic. To our surprise, without applying any exogenous stimulation, we observed intermittent calcium transients in the PHDs of *wildtype* animals, every 30 seconds on average (Fig. 6E, F and Movie 1). We observed calcium transients in 73% of the neurons imaged and no peaks could be identified in the remaining traces (n=19). Since in restrained animals there are small tail and spicule movements due to the defecation cycle and sporadic muscle contractions, one explanation could be that PHDs may be proprioceptive. We reasoned that if PHD activity resulted from internal tissue deformation caused by muscle contractions, inhibition of muscle activity should eliminate or reduce calcium transients. To inhibit muscle contractions, we generated transgenic worms in which muscles can be silenced in an inducible manner through the expression of the *Drosophila* histamine-gated chloride channel HisCl1 ^42^, under the *myo-3* promoter. To increase the efficiency of muscle silencing we introduced this transgene into an *unc-51(e369)* mutant background that renders animals lethargic. *unc-51* encodes a serine/threonine kinase required for axon guidance and is expressed in motor neurons and body wall muscle ^43^. Histamine-treated *myo-3::HisCl1;unc-51(e369)* animals were highly immobile and the frequency of calcium transients in the PHDs was strongly reduced (Fig. 6E, F and Movies 2 and S1). Transients were completely eliminated in half of the neurons (Fig. 6F). The reduction in transient frequency was specific to the silencing of the muscles because in histamine-treated *unc-51(e369)* and histamine non-treated *myo-3::HisCl1;unc-51(e369)* control animals, frequency was similar to that of *wildtype* animals (Fig. 6E, F and Movie S2).

Although PHDs are sensory neurons, they do receive some small synaptic input from other neurons (Fig. 6D) and therefore, the activity observed in restrained animals in response to muscle contractions could arise either directly through PHD sensory input or indirectly through presynaptic neurons. To test this, we imaged PHD activity in mutant males with impaired synaptic neurotransmission, *unc-13(e51)* ^44^ and dense core vesicle secretion, *unc-31(e169)* ^45^. In these mutants, PHD calcium transients persisted at similar frequencies to those in *wildtype* animals (Fig. 6E and F). This suggests that PHD activity does not require chemical input from the network. However, we cannot completely rule out that residual neurotransmission in the hypomorphic *unc-13(e51)* mutants may be sufficient to trigger wildtype levels of PHD activity in response to muscle contractions. Together, these results suggest that PHDs may respond directly to internal cues arising from muscle contractions, and that they may be proprioceptive neurons. PHDs may sense internal tissue deformations through their elaborate ciliated dendrites, which are deeply encased in the phasmid sheath and not exposed to the outside environment.

### A novel readjustment step during male-mating

The connectivity of PHD to male-specific neurons in the tail suggests that they may play a role in male reproductive behaviours controlled by tail circuits. In *C. elegans*, these include food-leaving behaviour, an exploratory strategy to search for mates ^46 47,48^, and mating behaviour. Mating consists in a temporal sequence of discrete behavioural steps: response (to mate contact); scanning (backward locomotion while scanning the mate’s body during vulva search); turning (at the end of the mate’s body to continue vulva search); location of vulva (stop at vulva); spicule insertion (intromission); and sperm transfer (reviewed in ^3^; Fig. 7). The coordinated execution of these mating step relies on sensory cues from the mate ^40^ and presumably, proprioceptive inputs within the male’s copulation circuit as well ^13^ (and reviewed in ^49^). This sensory information guides the male to either initiate the next step of the sequence or to reattempt the current, unsuccessful step through readjustment of movement and/or posture (reviewed in ^3^).

**Figure 7:**
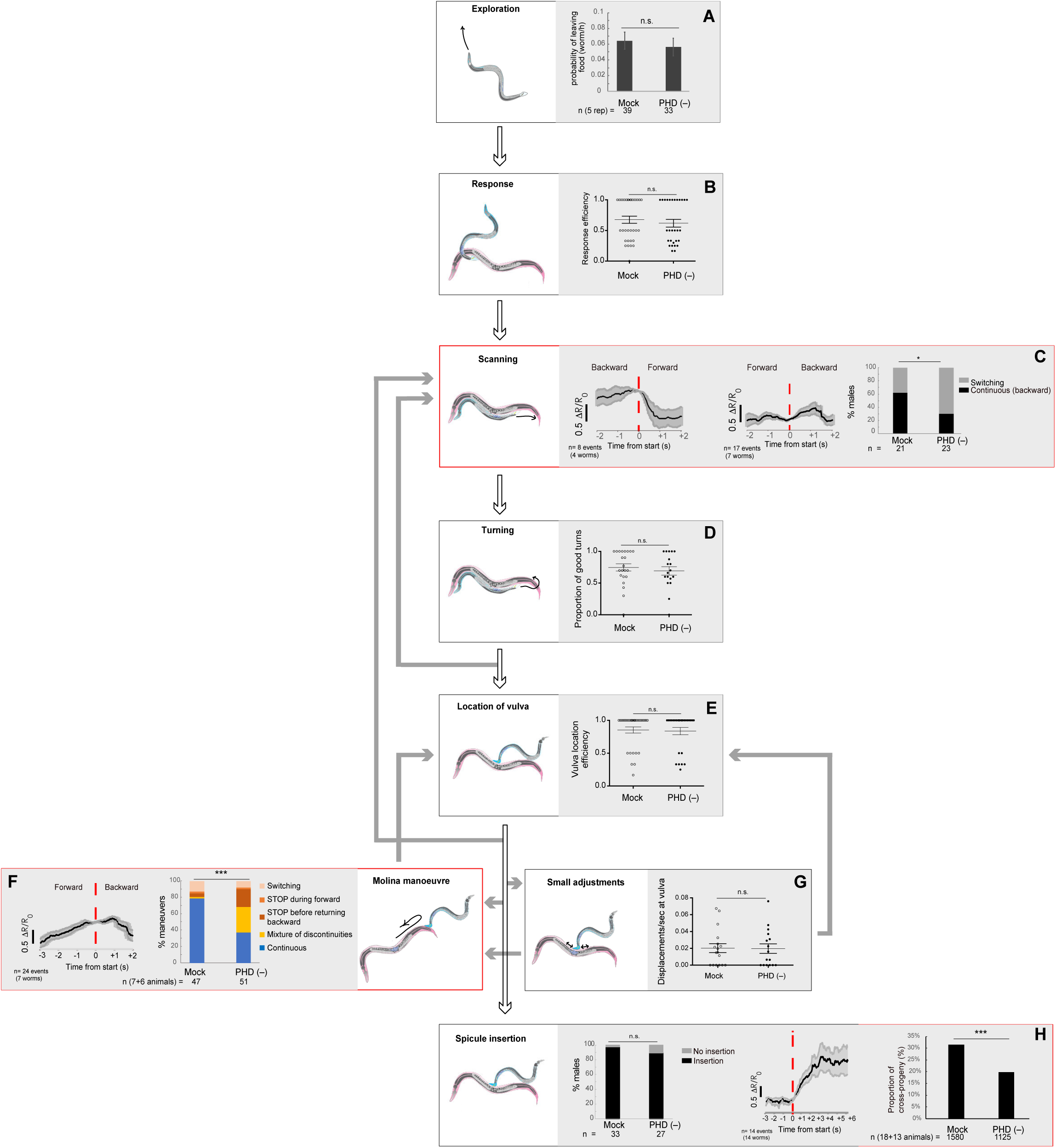
The PHD neurons are required for coordinated backward locomotion and effective intromission during mating. Diagram depicting the steps of reproductive behaviours controlled by the male tail circuits. The steps affected by PHD ablation are highlighted in red. Intact (mock) and ablated males carried either an *oig-8::gfp* or an *unc-17::gfp* transgene to identify the PHD neurons for ablation. White arrows indicate the transitions between the mating sequence. Grey arrows indicate the corrective transitions that males perform when they fail to attain the subsequent goal. The corrective transitions that occur upon failure of spicule insertion attempts (between location of vulva and spicule insertion) are always preceded by a displacement from the vulva (not depicted). Behavioural analysis in intact and PHD-ablated males are shown for each step. Calcium imaging in PHD neurons is shown for steps C, F and H. The black trace shows PHD activity as normalised GCaMP/RFP fluorescence ratio changes averaged for several events and phase locked (red dotted line) to the switch in the direction of locomotion (C and F) or to the start of spicule insertion (H). Grey shadow, S.E.M. A. Male exploratory behaviour measured as *P*_L_ values (probability of leaving food per worm per hour). n, number of males tested. Maximum likelihood statistical analysis was used to compare *P*_L_ values. n.s., no statistically significant difference, *P*≥0.05; error bars, S.E.M. B. Response efficiency, measured as 1/number of contacts with a mate before responding. C. Scanning locomotion during vulva search. *P*<0.05 (χ^2^ tested); n=animals test D. Turning, measured as proportion of good turns per male. E. Vulva location efficiency measured as 1/number of vulva encounters before stopping. F. Proportion of continuous and discontinuous Molina manoeuvres. Categories: switching (a forward movement away from the vulva interrupted repeatedly by a brief switch to backward movement without a STOP); STOP during forward (stopping during forward movement away from the vulva and then continuing in the same direction); STOP before returning backward (stopping at the transition between forward movement away from the vulva and returning backwards to the vulva); mixture (manoeuvres that displayed more than one of the discontinuities described above); continuous (smooth movement forward away from the vulva and return backwards to the vulva without stopping or switching in between). *P*<0.001 (χ^2^ test of continuous and discontinuous manoeuvres); n=number of events G. Number of displacements away from the vulva per unit of time spent at vulva. H. Left bar chart, proportion of males able to insert their spicules; n.s., no statistically significant difference, *P*≥0.05 (χ^2^ test); n= animals tested. Right bar chart, sperm transfer efficiency measured as percentage of cross-progeny; *P*<0.001 (χ^2^ test); n= total progeny. For B, D, E and G, bar and dots represent mean and individual animal values, respectively; error bars, S.E.M. n.s., no statistically significant difference, *P*≥0.05 (Mann-Whitney U test). Worm cartoons were modified with permission from original drawings by Rene Garcia.

During our behavioural analysis of *wildtype* males, we identified a novel readjustment movement that has not been previously described which we have termed the ‘*Molina manoeuvre’*. This movement occurs when the male has been trying to insert its spicules in the mate’s vulva for a period of time without success and subsequently loses vulva aposition. Unsuccessful spicule insertion attempts occurred in 65% of males, resulting in either a long displacement from the vulva, which led to the re-initiation of the scanning sequence, or in a small displacement (less than two tail-tip length) from the vulva (98% of vulva losses), as previously described ^50^. We find that these small displacements from the vulva lead to either local shifts to relocate the vulva (68% of total vulva losses) or in a Molina manoeuvre (32% of vulva losses) (Fig. 7). The Molina manoeuvre consists in the initiation of forward locomotion along the mate’s body away from the vulva to the end or middle of the mate’s body, at which point the male tail acquires a deep arched posture, followed by a return to the vulva with backward locomotion along the same route (Movie 3). Males perform this movement towards either end (head or tail) of the mate as a smooth, continuous sequence. Although we used paralysed *unc-51* mutant hermaphrodites to aid our mating analysis, males also perform this readjustment movement with moving *wildtype* mates (Movie S3).

### The PHD neurons are required for coordinated backward locomotion and effective intromission during mating

To test the hypothesis that PHDs regulate reproductive behaviours, we performed behavioural tests of intact and PHD-ablated males and functional imaging of PHD activity in freely behaving males during mating. We found no defects in food-leaving behaviour, response to mate contact, turning, location of vulva, or spicule insertion (Fig. 7A, B, D, E and H). However, these experiments revealed a role for PHDs in initiation and/or maintenance of backward locomotion during scanning and during the Molina manoeuvre. PHD-ablated males often switched their direction of locomotion during scanning, performing fewer continuous backward scans during vulva search compared to intact males (Fig. 7C). Consistent with this, we observed higher activity (i.e. Ca^2+^ levels) in PHD during backward locomotion than during forward locomotion while scanning (Fig. 7C). PHD activity also peaked just after the switch to backward locomotion during Molina manoeuvres, when the tail tip displays an acute bent posture, and PHD-ablated males performed defective, discontinuous manoeuvres, often stopping at the transition from forward to backward locomotion to return to the vulva (Fig. 7F and Movie 4). Importantly, we did not find significant differences between intact and PHD-ablated males in number of events that trigger the initiation of manoeuvres (Fig. 7G), indicating that PHDs are specifically required for coordinated locomotion during the manoeuvre itself, at the transition point to backward locomotion.Together, these data demonstrate that without intact PHD neurons, backward movement along the mating partner becomes slightly erratic, often interrupted by a switch in direction (during scanning), or its initiation is being delayed by a stop (during Molina manoeuvres). The qualitative difference in locomotion defects between these two steps of mating may result from differences in the state of the neural network before and after spicule insertion attempts ^50–52^.

In addition to the afore mentioned changes in neuronal activity, the highest level of PHD activation was observed during intromission, the penultimate step of the mating sequence, which precedes sperm transfer and involves full insertion of the spicules into the mate’s vulva while sustaining backward locomotion. PHD activity increased two-fold upon spicule insertion and remained high for several seconds while the spicules were inserted (Fig. 7H). PHDs are required for the efficiency of these two last steps of mating since PHD-ablated males that were observed to complete the mating sequence, including intromission and ejaculation, during a single mating with one individual mate, produced fewer cross-progeny than intact males in the same conditions (Fig. 7H). Intact PHDs may increase the efficiency of sperm transfer by controlling the male’s posture during intromission. While the cues and network interactions that activate PHDs during mating are presently not known, the high levels of PHD activity upon spicule insertion are consistent with a putative proprioceptive role for these neurons.

## Discussion

The data presented here extend John Sulston’s original observations and demonstrate that male PHso1 cells undergo a direct glia-to-neuron transdifferentiation to produce a novel class of bilateral cholinergic, peptidergic and ciliated sensory neurons, the phasmid D neurons. This updates the anatomy of the *C. elegans* male to be comprised of 387 neurons (93 of which are male-specific) and 90 glia. This is the second example of neurons arising from glia in *C. elegans*, confirming that glia can act as neural progenitors across metazoan taxa. In both cases we have demonstrated that fully differentiated, functional glia can retain neural progenitor properties during normal development. The complete glia-to-neuron cell fate switches we observe strongly suggest the production of neurons from glia is a process of natural transdifferentiation. In the first case, this is an indirect transdifferentiation as it occurs via an asymmetric cell division, leading to self-renewal of the glial cell and the production of a neuron ^14^. In the work described here it is a direct transdifferentiation and as such is the second direct transdifferentiation to be described in *C. elegans*. The first involves the transdifferentiation of the rectal epithelial cell Y into the motor neuron PDA, which occurs in hermaphrodites at earlier larval stages, before sexual maturation ^53^. It will be interesting to determine whether the mechanisms that regulate the PHso1-to-PHD cell fate switch are similar or distinct from those that have been described for Y-to-PDA ^54–56^. Intriguingly, direct conversion of glial-like neural stem cells has been observed in the adult zebrafish brain ^57^, suggesting that this may be a conserved mechanism for postembryonic generation of neurons.

Based on our manipulations and imaging of PHD activity in restrained and freely mating animals, we propose that PHDs may be proprioceptive neurons which become activated by tail tip deformations and engage the circuits for backward locomotion. The high levels of PHD activity during intromission and upon tail tip bending during the Molina manoeuvre, steps which require inhibition of forward locomotion and sustained backward movement, are consistent with such model. PHDs may ensure sustained backward locomotion through excitation of their postsynaptic targets PVY, PVX and EF interneurons, all of which have been shown to promote backward movement through AVA and AVB during scanning and location of vulva ^58,59^. As *wildtype* hermaphrodites are highly uncooperative during mating and actively move away ^60^, any erratic and uncoordinated movement by the male is likely to result in the loss of its mate.

Sensory input from the mating partner is essential for successful mating and accordingly, many sensory neurons in copulation circuits are dedicated to this purpose (reviewed in ^61^ and ^3^). However, coordinated motor control in all animals requires also proprioception, sensory feedback from internal tissues that inform the individual about its posture and strength exerted during movement ^62,63^. Several putative proprioceptive neurons have been identified in *C. elegans* (reviewed in ^64^). Within the male copulation circuit, only the spicule neuron SPC has been attributed a putative proprioceptive role, based on the attachment of its dendrite to the base of the spicules ^13^. The SPC neuron is required for full spicule protraction during intromission ^39,65^.The presence of proprioceptive neurons in copulation circuits may be a broadly employed mechanism to regulate behavioural transitions during mating. In this light, it would be interesting to determine whether the mechanosensory neurons found in mating circuits of several organisms ^66–68^ may play a role in self-sensory feedback.

In summary our results provide new insight into the developmental mechanisms underlying the neural substrates of sexually dimorphic behaviour. The glia-to-neuron transdifferentiation that results in PHD represents an extreme form of sexual dimorphism acquired by differentiated sex-shared glial cells during sexual maturation. The PHD neurons, perhaps through proprioception, enable the smooth and coordinated readjustment of the male’s movement along its mate when spicule insertion becomes difficult to attain. This readjustment movement represents an alternative step within the stereotyped mating sequence. These findings reveal the extent to which both cell fate and innate behaviour can be plastic yet developmentally wired.

## Materials and Methods

### Strains

For a list of strains generated and/or used in this study see Table S2

### DNA constructs and transgenic strains

The *oig-8::gfp* reporter was built by amplifying a 964 bp promoter fragment upstream of the *oig-8* ATG adding the restriction sites of SphI and XmaI. This PCR product was digested and ligated into SphI/XmaI-digested pPD95.75 vector. To generate *oig-8p::mCherry::rab-3*, the same *oig-8* promoter fragment was ligated into SphI/XmaI-digested *pkd-2p::mCherry::rab-3* (a gift from M. Lázaro-Peña). The *oig-8::GCaMP6f::SL2::RFP* construct was created by PCR fusion ^69^. A ~ 1 kb promoter region of the *oig-8* locus was amplified with primers: (A) gggagtgacctatgcaaacc and (B) CGACGTGATGAGTCGACCATtgttttacctgaaatctttt, which has a tail overlapping with the GCaMP sequence for the fusion PCR. The *myo-3::HisCl1::sl2::mCherry* construct was created as PCR fusion. A 2.2 kb promoter region of the *myo-3* locus was amplified with primers: (A) cgtgccatagttttacattcc and (B) gctagttgggctttgcatGCttctagatggatctagtggtc, which has a tail overlapping with the *HisCl1* sequence for the fusion PCR. The *lin-48::tdTomato* construct was a kind gift from Mike Boxem and contains a 6.8kb promoter fragment upstream of the coding sequence of *tdTomato* and the *unc-54* 3’ UTR. It was injected into *him-5(e1490)* animals and spontaneously integrated generating *drpIs3* (see Table S2).

The *grl-2::gfp* reporter was generated by integrating *sEx12852* ^26^.

### Cell ablations

PHD was ablated with a laser microbeam as previously described ^70^. L4 males were staged and placed in a seeded plate the night before. Ablations were carried out at 1 day adulthood and PHD was identified by *oig-8::gfp* or *unc-17::gfp* reporter expression. Mock-ablated animals underwent the same treatment as ablated males except for laser trigger. Animals were left to recover overnight to perform behavioural assays the next day. After behavioural assays, animals were checked for lack of GFP expression in PHD to confirm correct cell ablation. The few animals in which PHD had not been efficiently ablated were discarded from the data.

### Cell-type specific sex transformations

We used two previously described strains each containing an array that drives *fem-3* expression from a *grl-2* promoter fragment ^14^. In the case of *oleEx24* the presence/absence of the array in PHso1 and PHso2 was monitored by the *mCherry* expression from the array itself and the expression of an *ida-1::gfp* reporter (*inIs179*), in the background of the strain was used to monitor neuronal fate in PHso1/PHD. In the case of *oleEx18*, we had difficulty visualising the *mCherry* from the array in PHD and crossed a *lin-48::tdTomato* (*drpIs3*) into the strain to visualise PHso1/PHD. The presence/absence of the array in whole animals was assessed using the array co-injection marker *elt-2::gfp*. Neuronal fate was monitored using a *rab-3::yfp* reporter. See Table S2 for full strain details.

### Behavioural assays

All behavioural assays were scored blind to the manipulation. Males carried either an *oig-8::gfp* or an *unc-17::gfp* transgene to identify the PHD neurons for ablation.

#### Food leaving

Animals were tested at 3 days of adulthood (the day after they had been tested for mating). Assays were performed and scored as previously described ^47^.

#### Mating

Assays were performed and scored as previously described ^14^. Males were tested at 2 days of adulthood with 1 day-old *unc-51(e369)* hermaphrodites picked the night before as L4s. Each male was tested once for all steps of mating. Those males that were not successful at inserting their spicules with the first hermaphrodite were tested again with a maximum of three hermaphrodites to control for hermaphrodite-specific difficulty in penetration ^39^. Assays were replicated at least three times on different days and with different sets of males. Videos of matings were recorded and scored blind by two independent observers. Displacements from the vulva were scored as movements of one or two tail-tip distance away from the vulva.

#### Fertility assays

each individual male was monitored for all mating steps during a single mating until it ejaculated. After disengagement from the mate, the hermaphrodite was picked and placed in a fresh plate to lay progeny. The adult hermaphrodite was transferred to a fresh plate each day during three consecutive days. After three days from the eggs being laid, L4 larvae and adult progeny were counted as Unc self-progeny of Wt cross-progeny.

### Microcopy and imaging

Worms were anesthetized using 50mM sodium azide and on mounted on 5% agarose pads on glasss slides. Images were acquired on a Zeiss AxioImager using a Zeiss Colibri LED fluorescent light source and custom TimeToLive multichannel recording software (Caenotec). Representative images are shown following maximum intensity projections of 2-10 1*µ*m z-stack slices and was perfumed in ImageJ.

### Ca^2+^ imaging

Imaging was performed in an upright Zeiss Axio Imager 2 microscope with a 470nm LED and a GYR LED (CoolLED) with a dual-band excitation filter F59-019 and dichroic F58-019 (Chroma) in the microscope turret. Emission filters ET515/30M and ET641/75 and dichroic T565lprx-UF2 were placed in the cube of a Cairn OptoSplit II attached between the microscope and an ORCA-Flash 4 camera (Hamamatsu). Acquisition was performed at 20 fps.

Imaging during mating was performed with a 20× long working distance objective (LD Plan-NEOFLUAR numerical aperture 0.4), placing the male on an agar pad with food and 20 hermaphrodites. The ~ 50 mm per side agar pad was cut out from a regular, seeded NGM plate and placed on a glass slide. The hermaphrodites were placed in a ~ 100 mm^2^ centre region. A fresh pad was used every two recordings.

Imaging in restrained animals was performed for 2.5 to 3 minutes with a 63× objective (LD C-apochromat numerical aperture 1.15). Animals were glued with Wormglu along the body to a 5% agarose pad on a glass slide and covered with M9 or 20 mM histamine (Sigma, H7125) and a coverslip.

### Ca^2+^ imaging analysis

A moving region of interest in both channels was identified and mean fluorescent ratios (GFP/RFP) were calculated with custom-made Matlab scripts ^71^, kindly shared by Zoltan Soltesz and Mario de Bono. For recordings in restrained animals, bleach correction was applied to those traces in which an exponential decay curve fitted with an R square > 0.6. Ratios for each recording were smoothened using a 8 frame rolling average. For ΔR/R_0_ values, R_0_ for each recording period was calculated as the mean of the lowest 10^th^ percentile of ratio values. Traces that were locked to behavioural transitions had their ΔR/R_0_ values added or subtracted such that all events had the same value at t=0. Peaks were identified manually by an observer blind to the genotype and treatment. Peaks were called as signals above 0.2 ΔR/R_max_ (where R_max_ was calculated as the highest 5^th^ percentile of ratio values), above 2σ from local basal and a minimum duration of 5 seconds.

### Electron microscopy and serial reconstruction

The samples were fixed by chemical fixation or high-pressure freezing and freeze substitution as previously described ^72^. Several archival print series from *wildtype* male tails in the MRC/LMB collection were compared to *wildtype* adult males prepared in the Hall lab, showing the same features overall. Ultrathin sections were cut using a RMC Powertome XL, collected onto grids, and imaged using a Philips CM10 TEM. The PHD cell bodies were identified in the EM sections based on position and morphology. This was followed by serial tracing of the projections to establish their morphology and connectivity. The method of quantitative reconstruction using our custom software is described in detail in ^4,73^. The connectivity of the PHD neurons was determined from the legacy N2Y EM series ^13^. Circuit diagrams were generated using Cytoscape ^74^. PHD neuronal maps and connectivity tools are available at www.wormwiring.org.

## Author contributions

L.M.G. performed, analysed and interpreted the behavioural experiments; B.K. identified neuronal characteristics in PHD and generated the *oig-8* reporter constructs; S.J.C. with S.W.E reconstructed the connectivity; R.B. performed the sex-transformation experiments and with M.S. and J.M.O. performed the characterization of the PHso1-to-PHD molecular transdifferentiation; S.P.R.G scored *grl-*2 expression and histone condensation in larvae; D.J.E. provided technical assistance; D.H.H. provided and analysed the electron micrographs; A.B. conceived, performed and interpreted the behavioural and Ca^2+^ imaging experiments and co-wrote the manuscript; R.J.P. conceived and performed the analysis of the PHso1-to-PHD transdifferentiation and co-wrote the manuscript.

## Acknowledgements

We thank Mario de Bono and Zoltan Soltesz for kindly sharing custom-made software for ratiometric analysis of Ca^2+^ imaging in moving animals; Rene Garcia for worm cartoons; Ken Nguyen for help with EM; John G. White and Jonathan Hodgkin for their help in transferring archival TEM data from the MRC/LMB to the Hall Lab at the Albert Einstein College of Medicine for long-term curation and study; Shai Shaham for sharing the *mir-228::gfp* strain prior to publication; Mike Boxem, María Lázaro-Peña and Cori Bargmann for additional strains and reagents; Sheila Poole for edits on the manuscript; Baris Kuru for aid with behavioural experiments. Additional strains were obtained from the CGC, which is supported NIH grant <u>P40 OD010440</u>. Christopher Brittin was influential in designing and creating www.wormwiring.org. This work was supported by a Newton Fellowship from the Royal Society to L.M.G., a Wellcome Trust PhD studentship to R.B., Marie Curie CIG grant 618779 and Wellcome Trust Enhancement Funding (095722/Z/11/A) to R.J.P., NIH R01 GM066897 grant and the G. Harold & Leila Y. Mathers Charitable Foundation to S.W.E, NIH OD 010943 to D.H.H., NIH T32GM007491 and F32 MH115438 01 to S.J.C.; R.J.P. was supported by a Wellcome Trust Research Career Development Fellow (095722/Z/11/Z) and is currently a Wellcome Senior Fellow in Basic Biomedical Science (207483/Z/17/Z). A.B and R.J.P are members of COST Action BM1408.

## Movie Legends

Movies 1 and 2: Imaging of neuronal activity in PHD neurons with GCaMP6f (left channel) and RFP (right channel) in restrained animals. Animals are expressing a *oig-8::GCaMP6f::sl2::rfp* transgene. Movies play at 100 fps (recorded at 20 fps).

1. *wildtype* male
2. *unc-51(e359)* male expressing a histamine-inducible silencing transgene in muscle (*myo-3::HisCl1::mCherry*) and treated with 20 mM histamine. Movies 3 and 4: Males performing Molina manoeuvres during mating. Movies are played at 40 fps.
3. *wildtype* male performing a Molina manoeuvre during mating with a paralysed *unc-51(e359)* hermaphrodite.
4. PHD-ablated male performing a defective, discontinuous Molina manoeuvre.

## References

1. Yang, C. F. & Shah, N. M. Representing sex in the brain, one module at a time. Neuron 82, 261–278 (2014).

2. Auer, T. O. & Benton, R. Sexual circuitry in Drosophila. Curr Opin Neurobiol 38, 18–26 (2016).

3. Barr, M. M., Garcia, L. R. & Portman, D. S. Sexual Dimorphism and Sex Differences in Caenorhabditis elegans Neuronal Development and Behavior. Genetics 208, 909–935 (2018).

4. Jarrell, T. A. et al. The Connectome of a Decision-Making Neural Network. Science (New York, NY) 337, 437–444 (2012).

5. Ryan, D. A. et al. Sex, age, and hunger regulate behavioral prioritization through dynamic modulation of chemoreceptor expression. Curr. Biol. 24, 2509–2517 (2014).

6. Oren-Suissa, M., Bayer, E. A. & Hobert, O. Sex-specific pruning of neuronal synapses in Caenorhabditis elegans. Nature 533, 206–211 (2016).

7. Hilbert, Z. A. & Kim, D. H. Sexually dimorphic control of gene expression in sensory neurons regulates decision-making behavior inC. elegans. Elife 6, 386 (2017).

8. Serrano-Saiz, E. et al. A Neurotransmitter Atlas of the Caenorhabditis elegans Male Nervous System Reveals Sexually Dimorphic Neurotransmitter Usage. Genetics 206, 1251–1269 (2017).

9. Serrano-Saiz, E., Oren-Suissa, M., Bayer, E. A. & Hobert, O. Sexually Dimorphic Differentiation of a C. elegans Hub Neuron Is Cell Autonomously Controlled by a Conserved Transcription Factor. Curr. Biol. 27, 199–209 (2017).

10. Weinberg, P., Berkseth, M., Zarkower, D. & Hobert, O. Sexually Dimorphic unc-6/Netrin Expression Controls Sex-Specific Maintenance of Synaptic Connectivity. Curr. Biol. 28, 623–629.e3 (2018).

11. Pereira, L. et al. Timing mechanism of sexually dimorphic nervous system differentiation. Elife 8, 2467 (2019).

12. Sulston, J. E. & Horvitz, H. R. Post-embryonic cell lineages of the nematode, Caenorhabditis elegans. Dev Biol 56, 110–156 (1977).

13. Sulston, J. E., Albertson, D. G. & Thomson, J. N. The Caenorhabditis elegans male: postembryonic development of nongonadal structures. Dev Biol 78, 542–576 (1980).

14. Sammut, M. et al. Glia-derived neurons are required for sex-specific learning in C. elegans. Nature 526, 385–390 (2015).

15. Emmons, S. W. Neural Circuits of Sexual Behavior in Caenorhabditis elegans. Annu. Rev. Neurosci. 41, 349–369 (2018).

16. Sulston, J. E., Schierenberg, E., White, J. G. & Thomson, J. N. The embryonic cell lineage of the nematode Caenorhabditis elegans. Dev Biol 100, 64–119 (1983).

17. Ward, S., Thomson, N., White, J. G. & Brenner, S. Electron microscopical reconstruction of the anterior sensory anatomy of the nematode Caenorhabditis elegans. J. Comp. Neurol. 160, 313–337 (1975).

18. White, J. G., Southgate, E., Thomson, J. N. & Brenner, S. The structure of the nervous system of the nematode Caenorhabditis elegans. Philos Trans R Soc Lond B Biol Sci 314, 1–340 (1986).

19. Bird, A. F. & Bird, J. The Structure of Nematodes. (Academic Press, 1991).

20. Doroquez, D. B., Berciu, C., Anderson, J. R., Sengupta, P. & Nicastro, D. A high-resolution morphological and ultrastructural map of anterior sensory cilia and glia in Caenorhabditis elegans. Elife 3, e01948 (2014).

21. Low, I. I. C. et al. Morphogenesis of neurons and glia within an epithelium. Development 146, dev171124 (2019).

22. Perkins, L. A., Hedgecock, E. M., Thomson, J. N. & Culotti, J. G. Mutant sensory cilia in the nematode Caenorhabditis elegans. Dev Biol 117, 456–487 (1986).

23. Johnson, A. D., Fitzsimmons, D., Hagman, J. & Chamberlin, H. M. EGL-38 Pax regulates the ovo-related gene lin-48 during Caenorhabditis elegans organ development. Development 128, 2857–2865 (2001).

24. Wildwater, M., Sander, N., de Vreede, G. & van den Heuvel, S. Cell shape and Wnt signaling redundantly control the division axis of C. elegans epithelial stem cells. Development 138, 4375–4385 (2011).

25. Wallace, S. W., Singhvi, A., Liang, Y., Lu, Y. & Shaham, S. PROS-1/Prospero Is a Major Regulator of the Glia-Specific Secretome Controlling Sensory-Neuron Shape and Function in C. elegans. Cell Rep 15, 550–562 (2016).

26. Hao, L., Johnsen, R., Lauter, G., Baillie, D. & Bürglin, T. R. Comprehensive analysis of gene expression patterns of hedgehog-related genes. BMC Genomics 7, 280 (2006).

27. Stefanakis, N., Carrera, I. & Hobert, O. Regulatory Logic of Pan-Neuronal Gene Expression in C. elegans. Neuron 87, 733–750 (2015).

28. Alfonso, A., Grundahl, K., Duerr, J. S., Han, H. P. & Rand, J. B. The Caenorhabditis elegans unc-17 gene: a putative vesicular acetylcholine transporter. Science (New York, NY) 261, 617–619 (1993).

29. Pereira, L. et al. A cellular and regulatory map of the cholinergic nervous system of C.elegans. Elife 4, e12432 (2015).

30. Zahn, T. R., Macmorris, M. A., Dong, W., Day, R. & Hutton, J. C. IDA-1, a Caenorhabditis elegans homolog of the diabetic autoantigens IA-2 and phogrin, is expressed in peptidergic neurons in the worm. J. Comp. Neurol. 429, 127–143 (2001).

31. Collet, J., Spike, C. A., Lundquist, E. A., Shaw, J. E. & Herman, R. K. Analysis of osm-6, a gene that affects sensory cilium structure and sensory neuron function in Caenorhabditis elegans. Genetics 148, 187–200 (1998).

32. Howell, K. & Hobert, O. Morphological Diversity of C. elegans Sensory Cilia Instructed by the Differential Expression of an Immunoglobulin Domain Protein. Curr. Biol. 27, 1782–1790.e5 (2017).

33. White, J. Q. et al. The sensory circuitry for sexual attraction in C. elegans males. Current Biology 17, 1847–1857 (2007).

34. Lee, K. & Portman, D. S. Neural Sex Modifies the Function of a C. elegans Sensory Circuit. Current Biology 17, 1858–1863 (2007).

35. White, J. Q. & Jorgensen, E. M. Sensation in a single neuron pair represses male behavior in hermaphrodites. Neuron 75, 593–600 (2012).

36. Fagan, K. A. et al. A Single-Neuron Chemosensory Switch Determines the Valence of a Sexually Dimorphic Sensory Behavior. Current Biology 1–27 (2018). doi:10.1016/j.cub.2018.02.029

37. Hodgkin, J. A genetic analysis of the sex-determining gene, tra-1, in the nematode Caenorhabditis elegans. Genes Dev 1, 731–745 (1987).

38. Zarkower, D. Somatic sex determination. WormBook 1–12 (2006). doi:10.1895/wormbook.1.84.1

39. Liu, K. S. & Sternberg, P. W. Sensory regulation of male mating behavior in Caenorhabditis elegans. Neuron 14, 79–89 (1995).

40. Barr, M. M. & Sternberg, P. W. A polycystic kidney-disease gene homologue required for male mating behaviour in C. elegans. Nature 401, 386–389 (1999).

41. Koo, P. K., Bian, X., Sherlekar, A. L., Bunkers, M. R. & Lints, R. The robustness of Caenorhabditis elegans male mating behavior depends on the distributed properties of ray sensory neurons and their output through core and male-specific targets. Journal of Neuroscience 31, 7497–7510 (2011).

42. Pokala, N., Liu, Q., Gordus, A. & Bargmann, C. I. Inducible and titratable silencing of Caenorhabditis elegans neurons in vivo with histamine-gated chloride channels. Proceedings of the National Academy of Sciences 111, 2770–2775 (2014).

43. Ogura, K., Shirakawa, M., Barnes, T. M., Hekimi, S. & Ohshima, Y. The UNC-14 protein required for axonal elongation and guidance in Caenorhabditis elegans interacts with the serine/threonine kinase UNC-51. Genes Dev 11, 1801–1811 (1997).

44. Miller, K. G. et al. A genetic selection for Caenorhabditis elegans synaptic transmission mutants. Proc Natl Acad Sci U S A 93, 12593–12598 (1996).

45. Speese, S. et al. UNC-31 (CAPS) is required for dense-core vesicle but not synaptic vesicle exocytosis in Caenorhabditis elegans. Journal of Neuroscience 27, 6150–6162 (2007).

46. Lipton, J., Kleemann, G., Ghosh, R., Lints, R. & Emmons, S. W. Mate searching in Caenorhabditis elegans: a genetic model for sex drive in a simple invertebrate. Journal of Neuroscience 24, 7427–7434 (2004).

47. Barrios, A., Nurrish, S. & Emmons, S. W. Sensory regulation of C. elegans male mate-searching behavior. Curr. Biol. 18, 1865–1871 (2008).

48. Barrios, A., Ghosh, R., Fang, C., Emmons, S. W. & Barr, M. M. PDF-1 neuropeptide signaling modulates a neural circuit for mate-searching behavior in C. elegans. Nat Neurosci 15, 1675–7682 (2012).

49. Garcia, L. R. Regulation of sensory motor circuits used in C. elegans male intromission behavior. Semin Cell Dev Biol 33, 42–49 (2014).

50. Correa, P., LeBoeuf, B. & Garcia, L. R. C. elegans dopaminergic D2-like receptors delimit recurrent cholinergic-mediated motor programs during a goal-oriented behavior. PLoS Genet. 8, e1003015 (2012).

51. Liu, Y. et al. A cholinergic-regulated circuit coordinates the maintenance and bi-stable states of a sensory-motor behavior during Caenorhabditis elegans male copulation. PLoS Genet. 7, e1001326 (2011).

52. Correa, P. A., Gruninger, T. & Garcia, L. R. DOP-2 D2-Like Receptor Regulates UNC-7 Innexins to Attenuate Recurrent Sensory Motor Neurons during C. elegans Copulation. Journal of Neuroscience 35, 9990–10004 (2015).

53. Jarriault, S., Schwab, Y. & Greenwald, I. A Caenorhabditis elegans model for epithelial-neuronal transdifferentiation. Proceedings of the National Academy of Sciences 105, 3790–3795 (2008).

54. Richard, J. P. et al. Direct in vivo cellular reprogramming involves transition through discrete, non-pluripotent steps. Development 138, 1483–1492 (2011).

55. Kagias, K., Ahier, A., Fischer, N. & Jarriault, S. Members of the NODE (Nanog and Oct4-associated deacetylase) complex and SOX-2 promote the initiation of a natural cellular reprogramming event in vivo. Proceedings of the National Academy of Sciences 109, 6596–6601 (2012).

56. Zuryn, S. et al. Transdifferentiation. Sequential histone-modifying activities determine the robustness of transdifferentiation. Science (New York, NY) 345, 826–829 (2014).

57. Barbosa, J. S. et al. Neurodevelopment. Live imaging of adult neural stem cell behavior in the intact and injured zebrafish brain. Science (New York, NY) 348, 789–793 (2015).

58. Sherlekar, A. L. et al. The C. elegans male exercises directional control during mating through cholinergic regulation of sex-shared command interneurons. PLoS ONE 8, e60597 (2013).

59. Sherlekar, A. L. & Lints, R. Nematode Tango Milonguero - the C. elegans male’s search for the hermaphrodite vulva. Semin Cell Dev Biol 33, 34–41 (2014).

60. Kleemann, G. A. & Basolo, A. L. Facultative decrease in mating resistance in hermaphroditic Caenorhabditis elegans with self-sperm depletion. 74, 1339–1347 (2007).

61. Villella, A. & Hall, J. C. Neurogenetics of courtship and mating in Drosophila. Adv. Genet. 62, 67–184 (2008).

62. Sherrington, C. S. On the proprioceptive system, especially in its reflex. Brain 29, 467–482 (1907).

63. Tuthill, J. C. & Azim, E. Proprioception. Curr. Biol. 28, R194–R203 (2018).

64. Schafer, W. R. Mechanosensory molecules and circuits in C. elegans. Pflugers Arch. 467, 39–48 (2015).

65. Garcia, L. R., Mehta, P. & Sternberg, P. W. Regulation of distinct muscle behaviors controls the C. elegans male’s copulatory spicules during mating. Cell 107, 777–788 (2001).

66. Liu, T. et al. FMRFamide-like neuropeptides and mechanosensory touch receptor neurons regulate male sexual turning behavior in Caenorhabditis elegans. Journal of Neuroscience 27, 7174–7182 (2007).

67. Ng, C. S. & Kopp, A. Sex combs are important for male mating success in Drosophila melanogaster. Behav. Genet. 38, 195–201 (2008).

68. Pavlou, H. J. et al. Neural circuitry coordinating male copulation. Elife 5, 3095 (2016).

69. Hobert, O. PCR fusion-based approach to create reporter gene constructs for expression analysis in transgenic C. elegans. Biotech. 32, 728–730 (2002).

70. Bargmann, C. I. & Avery, L. Laser killing of cells in Caenorhabditis elegans. Methods Cell Biol 48, 225–250 (1995).

71. Busch, K. E. et al. Tonic signaling from O₂ sensors sets neural circuit activity and behavioral state. Nat Neurosci 15, 581–591 (2012).

72. Hall, D. H., Hartwieg, E. & Nguyen, K. C. Q. Modern electron microscopy methods for C. elegans. Methods Cell Biol 107, 93–149 (2012).

73. Xu, M. et al. Computer assisted assembly of connectomes from electron micrographs: application to Caenorhabditis elegans. PLoS ONE 8, e54050 (2013).

74. Smoot, M. E., Ono, K., Ruscheinski, J., Wang, P.-L. & Ideker, T. Cytoscape 2.8: new features for data integration and network visualization. Bioinformatics 27, 431–432 (2011).

