## Supplementary Figure and Legends for "A direct glia-to-neuron natural transdifferentiation ensures nimble sensory-motor coordination of male mating behaviour"

**Supplementary Figure Legend**

**Figure S1: Ultrastructure of the male phasmid sensilla**

Transmission electron microscopy images from serial transverse thin sections of an adult male tail, from anterior (A) to posterior (E). Dorsal is up. The level at which the sections are taken are labelled on the figure according to the cilia structures visible in PHD (compare to Fig. S1). In all sections the left and right PHD neurons are indicated with a boxed red R(ight) or L(eft). In A, the other phasmid neurons are also labelled and their cilia can be observed in each panel.

A. The basal body of the cilia is visible in PHDL and PHDR.
B. The PHD cilia are first visible in this section.
C. The PHD axonemes are visible in this section.
D. The finger-like villi are labelled in this section with arrowheads (but are also visible in all sections), and can be traced in serial sections to the basal body of PHD. Their appearance is identical to the more numerous villi that extend from the AFD cilium in the amphid [1].

**Supplemental Movies:**

Movies S1 and S2: Imaging of neuronal activity in PHD neurons with GCaMP6f (left channel) and RFP (right channel) in restrained animals. Animals are expressing a *oig-8::GCaMP6f::sl2::rfp* transgene. Movies play at 100 fps (recorded at 20 fps).

S1. *unc-51(e359)* male expressing a histamine-inducible silencing transgene in muscle (*myo-3::HisCl1::mCherry*) and treated with 20 mM histamine.
S2. *unc-51(e359)* male treated with 20 mM histamine.

Movie S3: *wildtype* male performing a Molina manoeuvre during mating with a *wildtype* hermaphrodite. Movie is played at 40 fps.

1. Ward, S., Thomson, N., White, J. G. & Brenner, S. Electron microscopical reconstruction of the anterior sensory anatomy of the nematode Caenorhabditis elegans. *J. Comp. Neurol.* **160,** 313–337 (1975).

**Figure S1**

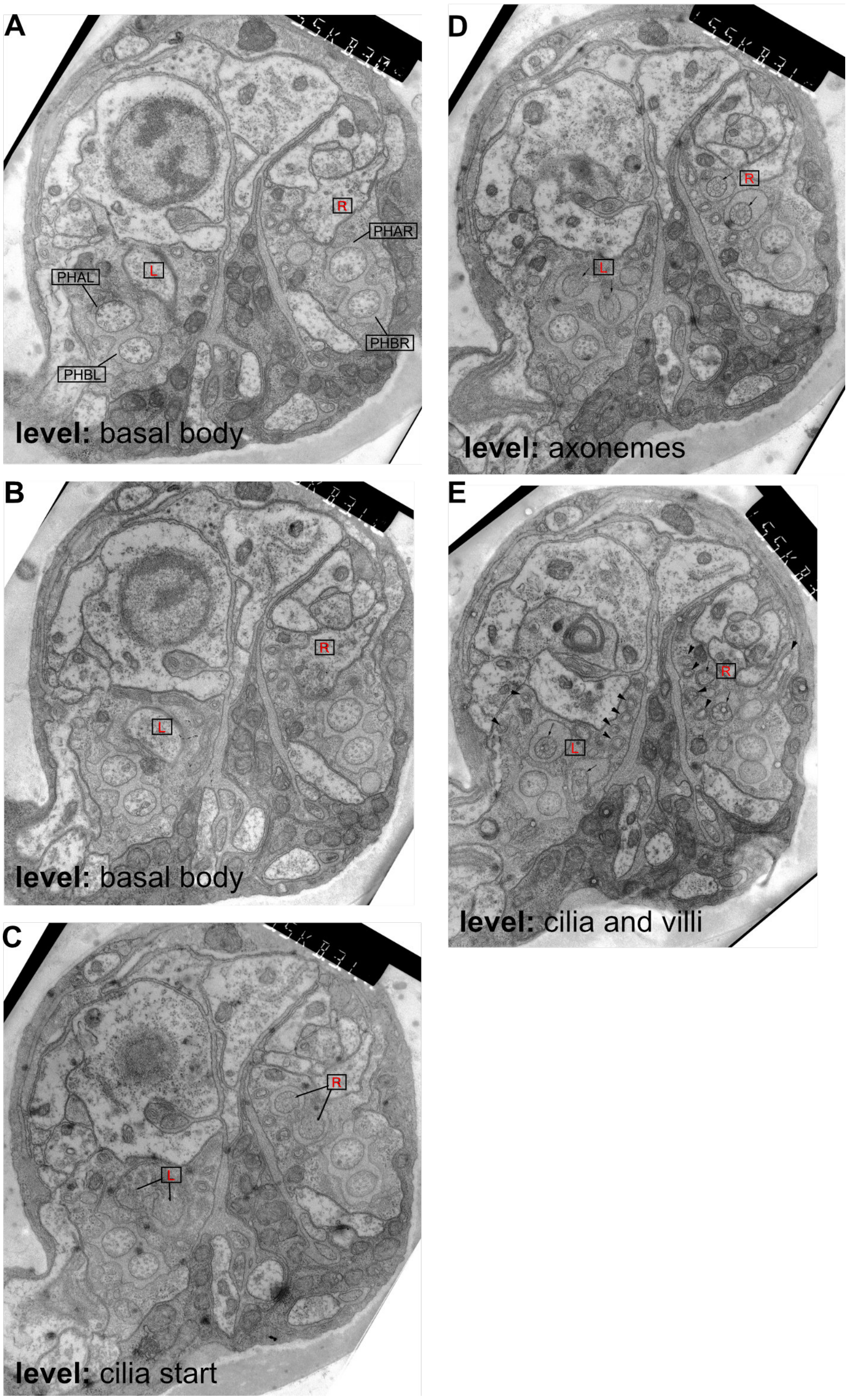
A
R
PHAR
L
PHAL
PHBR
PHBL
level: basal body
B
R
L
level: basal body
C
R
L
level: cilia start
D
R
L
level: axonemes
E
R
L
level: cilia and villi
